# The contribution of genetic variation of *Streptococcus pneumoniae* to the clinical manifestation of invasive pneumococcal disease

**DOI:** 10.1101/169722

**Authors:** Amelieke JH Cremers, Fredrick M Mobegi, Christa van der Gaast –de Jongh, Michelle van Weert, Fred J van Opzeeland, Minna Vehkala, Mirjam J Knol, Hester J Bootsma, Niko Välimäki, Jacques F Meis, Stephen Bentley, Sacha AFT van Hijum, Jukka Corander, Aldert L Zomer, Gerben Ferwerda, Marien I de Jonge

## Abstract

**Background:** Different clinical manifestations of invasive pneumococcal disease (IPD) have thus far mainly been explained by patient characteristics. Here we studied the contribution of pneumococcal genetic variation to IPD phenotype.

**Methods:** The index cohort consisted of 349 patients admitted to two Dutch hospitals between 2000-2011 with pneumococcal bacteraemia. We performed genome-wide association studies to identify pneumococcal lineages, genes and allelic variants associated with 23 clinical IPD phenotypes. The identified associations were validated in a nationwide (n=482) and a post-pneumococcal vaccination cohort (n=121). The contribution of confirmed pneumococcal genotypes to the clinical IPD phenotype, relative to known clinical predictors, was tested by regression analysis.

**Findings:** The presence of pneumococcal gene *slaA* was a nationwide confirmed independent predictor of meningitis (OR=10.5, p=0.001), as was sequence cluster 9 (OR=3.68, p=0.057). A set of 4 pneumococcal genes co-located on a prophage was a confirmed independent predictor of 30-day mortality (OR=3.4, p=0.003). We could detect the pneumococcal variants of concern in these patients’ blood samples by molecular amplification. In the post-vaccination cohort where the distribution of both patient characteristics and pneumococcal serotypes had changed, the relative importance of the prophage was no longer supported.

**Interpretation:** Knowledge of pneumococcal genotypic variants improved our clinical risk assessment for detrimental manifestations of IPD. This provides us with novel opportunities to target, anticipate or avert the pathogenic effects that are related to particular pneumococcal variants. Therefore, future diagnostics should facilitate prompt appreciation of pathogen diversity in clinical sepsis management. Ongoing surveillance is warranted to monitor the clinical value of information on pathogen variants in dynamic microbial and susceptible host populations.

**Funding:** None.

## Background

Invasive pneumococcal disease (IPD) is a threat to both the patient as well as the pneumococcus.^1^ It occurs nonetheless, and is a major cause of morbidity and mortality worldwide.^2^ The variety in clinical presentations across IPD patients is considerable, and not fully explained by host factors alone.^3^ It is therefore of interest to investigate whether it matters which pneumococcal variant happens to proliferate in the body.

Invasive disease includes ongoing presence of bacteria in blood and further sterile body sites like the pleural cavity and cerebrospinal fluid, corresponding with the clinical syndromes bacteraemia, empyema, and meningitis respectively. The reasons for these phenomena to occur mainly include flaws in host defence.^4-6^ Although “invasive” pneumococcal traits have been suggested as well,^7,8^ the large variety of pneumococci retrieved from IPD and the replacement of serotypes observed after the introduction of pneumococcal conjugate vaccines (PCVs) temper the importance of pneumococcal variation as determinant of invasive disease.^9,10^

Patients who have acquired pneumococci in their bloodstream do not always develop sepsis and clinical presentations vary from mild respiratory disease to imminent death.^11^ Aside from the classical vulnerable elderly patient who slowly recovers from pneumococcal pneumonia upon in-hospital treatment, IPD can manifest at all ages, in a range of body sites, with varying severity and sequelae. It is important to understand the origins of this diversity. Despite the introduction of uniform clinical guidelines and vaccines, the global pneumococcal disease burden remains high ^9,12,13^ and patients may benefit from more tailored adjunctive measures targeting the effects of specific pneumococcal variants. ^14^

The diversity in pneumococcal variants, illustrated by over 95 different capsular serotypes, has long been appreciated in pneumococcal vaccination and surveillance. *S. pneumoniae* is a naturally competent organism that fosters genetic recombination via transformation throughout its entire genome.^15^ Although pneumococcal serotypes have been related to particular clinical manifestations of IPD,^16^ it is unsure if the capsule is solely responsible.

Here we studied whether genome-wide pneumococcal variants were associated with clinical manifestations of human IPD in naturally occurring patient populations.

## Methods

### Three clinical cohorts

The index cohort consisted of 349 patients diagnosed with a pneumococcal bacteraemia admitted to two Dutch hospitals between January 2000 and June 2011. For the geographical validation cohort 482 adults with IPD admitted to 20 other Dutch hospitals (having blood cultures assessed in 9 sentinel laboratories) between June 2004 - December 2006 and June 2008 - May 2012 (periods for which clinical metadata were available) were randomly selected from the National Surveillance Database.^17,18^ The temporal validation cohort was collected from one index hospital, and consisted of 121 pneumococcal bacteraemia patients hospitalized between November 2012 and February 2016. In the latter cohort the distribution of pneumococcal serotypes had markedly changed since the introduction of PCVs in the Dutch National Immunisation Programme for infants (7-valent PCV in 2006, 10-valent PCV in 2011). This observational study was approved by the Medical Ethical Committees of the participating hospitals. Clinical data were collected from medical charts. Details on handling and formatting of clinical variables for each analysis are described in Supplementary methods 1. Normality of continuous variables was tested by Shapiro-Wilk test, and differences in characteristics in comparison to the index cohort were tested with a 2-sided Student’s t-test or Mann-Whitney U test accordingly. Differences in nominal variables were tested by 2-sided Chi-square testing (Fisher’s exact if less than 10 cases in any cell). Pneumococcal blood isolates were stored in 10% glycerol 10% skim milk at −80°C. Serotypes were determined by capsular PCR and Quellung reaction, confirmed by molecular capsular typing in whole genome sequenced strains. From isolates at the index hospitals, DNA was isolated with Qiagen Genomic-tip 20/G after culture to OD_620_ 0.2-0.3 in 10ml Todd Hewitt broth with 5% yeast extract at 37°C. DNA template from the National Surveillance isolates was prepared as a lysate by heating a 2ml overnight culture at 90°C for 10 minutes.

### Genome-wide analyses in the index cohort

Whole genome sequencing and assembly, as well as determination of orthologous genes (OGs), functional annotations, core genome, population phylogeny, and population structure (i.e. the identification of genetically diverged subpopulations which are called sequence clusters) were performed for the 349 isolates from the index cohort as previously described.^10^

The relationship between sequence cluster (SC) and 23 clinical IPD phenotypes was explored by stepwise regression analysis. Dependent variables were clinical IPD phenotypes, and each sequence cluster was entered as a binary independent variable. The models included a constant, with variable entry set at 0.05, and removal at 0.05 for logistic and at 0.1 for linear regression.

We performed two different genome wide association studies in the index cohort. First we investigated the relationship between the presence of each individual OG on the accessory pneumococcal genome and the clinical IPD phenotype. For this analysis OGs present in <98% and >2% of cases were selected. Associations with binary clinical variables were assessed by Fisher’s exact with cluster permutation and by Cochran-Mantel-Haenszel analyses implemented in PLINK.^19^ Sequence cluster (SC) was introduced as the nominal covariate to adjust for population structure, and p-values were false discovery rate-corrected to adjust for multiple testing by the Benjamini-Hochberg procedure. Second we investigated the relationship between any allelic variant present anywhere on the genome and the clinical IPD phenotype. K-mers (DNA-words of 10 to 99 base pairs) were identified from draft assemblies by distributed string mining, and subsequently filtered for adjacent bases having a different frequency support vector in the study cohort, and for being associated with each phenotype at p-values < 1e-5 in univariate chi-square testing. Associations between the selected k-mers and binary clinical IPD phenotypes were assessed by sequence element enrichment (SEER) analysis,^20^ including correction for population structure by multi-dimensional scaling using a random subset of k-mers. The origin of k-mers was determined by alignment to the annotated draft genomes of the index cohort with complete coverage and identity using BLAST. To adjust for multiple testing, the significance threshold was set at 1e-8.

### Validation of associations

We aimed to validate a selection of the identified OGs that were significantly associated with a clinical IPD phenotype, irrespective of their functional annotation, in a nation-wide cohort. In the temporal validation cohort the number of identified genes evaluated was constrained by the number of cases in that collection period. The size of the validation cohorts was calculated to detect the index differences with a power of at least 0.8 and alpha of 0.05 in a 1-sided fashion. Because the similarity in distribution of phenotypes and OGs in the validation cohorts was uncertain, the significance threshold for validation was set at 0.1.

To determine the presence of the OGs of interest on pneumococcal genomes in the validation cohorts, primers were designed and validated based on the index cohort, using a real-time fluorescent read out. The 20μl reaction mix contained 1x SsoAdvanced universal SYBR Green supermix, 200nM of each primer and as template either 0.005ng Qiagen genomic tip DNA or 200 times diluted pneumococcal lysate which yielded similar Ct-values. Cycling conditions were 95°C 3min; 40 cycles of 95°C 10sec and 55°C 30sec; 95°C 10sec; melting curve 65 to 95°C with 0.5°C/sec increase. All diluted templates were tested for detection of the *gyrA* pneumococcal housekeeping gene. All PCR runs included positive and negative control samples from the index cohort, plus negative extraction and PCR controls, and the specificity of produced amplicon for the OG of interest was confirmed by its melting temperature (Supplementary methods 2).

### Confirmed pneumococcal genotypes

Co-occurrence of confirmed genotypes with other sequence variants was determined by Pearson correlation. Co-localization of confirmed OGs with bacteriophages was assessed by identification of predicted prophage sequences in the draft genomes of the index cohort using PHASTER.^21^ Sequence variation within the confirmed OGs and prophages was expressed in size, GC-content and pairwise distances. Distances were calculated from amino acid alignments, using the MEGA7 p-distance metric assuming gamma distribution with pairwise deletion of ambiguous positions.

### Clinical relevance

The relative contribution of the identified pneumococcal genotypes to the clinical IPD phenotype in relation to well-known clinical predictors was assessed by logistic regression analysis as described elsewhere,^22^ with the addition of the significantly associated sequence clusters.

To explore clinical detection of pneumococcal variants during IPD, stored serum samples collected from IPD patients at day 0-3 of hospitalization were retrieved from −40°C. The pneumococcal genomic DNA load in the serum samples was assessed previously.^23^ We selected those serum samples on which capsular sequence typing had previously been successful.^24^ DNA was isolated from 100μl of serum using Qiagen’s DNeasy Blood and tissue kit. The OG validation PCRs (not matching human DNA sequences) were performed in duplicate using 8μl of template DNA and 50 amplification cycles.

Unless stated otherwise, the significance threshold was set at 0.05.

## Results

### Three clinical cohorts

Although the geographical validation cohort largely overlapped with the study period of the index cohort, serotypes appeared to be not evenly distributed (Figure 1). The temporal validation cohort was included to monitor identified associations in changing populations. In this cohort the patient characteristics of IPD cases had altered as compared to the index cohort, and serotypes clearly changed in response to pneumococcal vaccination. However, the distribution of IPD syndromes and outcomes had remained stable over time.

**Figure 1.**
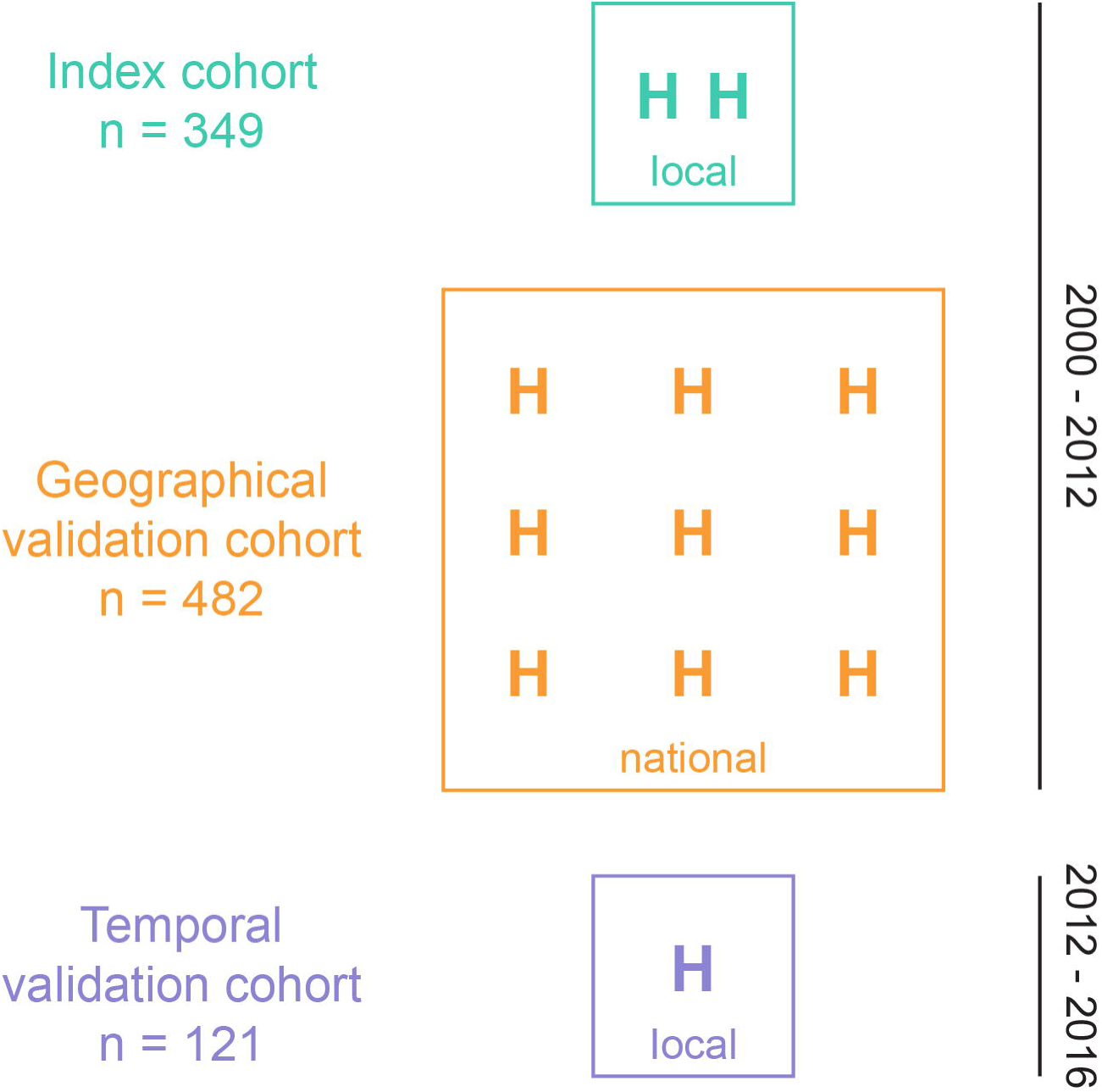
Cohort characteristics. The index cohort consisted of 349 patients with a pneumococcal bacteraemia admitted to 2 local hospitals (H), the geographical validation cohort of 482 patients in nationwide IPD surveillance, and the temporal validation cohort of 121 patients admitted to the index hospitals during a later time period (panel A). Main cohort characteristics are presented as mean ± standard deviation, median (interquartile range), or percentage fulfilling the condition (N/N known) (panel B). ^∗^:p<0.05; ^∗∗^:p<0.01; ^∗∗∗^:p<0.001. *Abbreviations:* PSI: pneumonia severity index; PCV7: serotypes 4, 6B, 9V, 14, 18, and 23F; PCV10: serotypes 1, 5, and 7F; PCV13: serotypes 3, 6A, and 19A; NVT: all other (non vaccine) serotypes. P-values support differences from the index cohort.

### Genome-wide analyses in the index cohort

Of the 23 tested clinical manifestations of IPD 87% appeared to be associated with one or more pneumococcal sequence clusters (SCs) (Table 1).

**Table 1.**
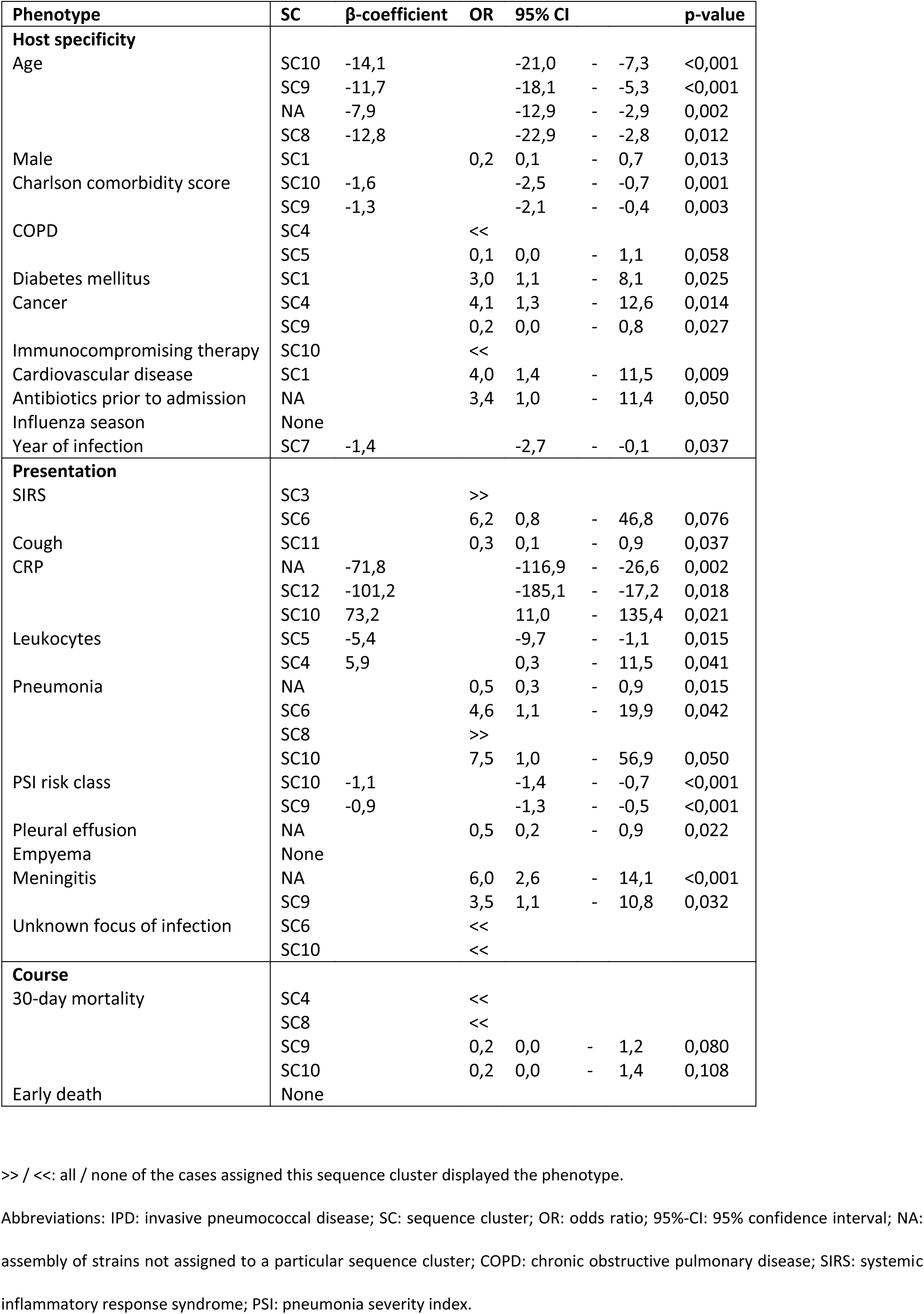
Sequence cluster as determinant of clinical IPD phenotype in the index cohort

In the first GWAS we studied the relationship between the presence of individual orthologous genes (OGs) on the accessory pneumococcal genome and the clinical IPD phenotype. Independently from SC, 68 of the 1127 selected pneumococcal OGs were associated with nine different clinical IPD phenotypes (Supplementary file 1, and most pronounced associations displayed in Figure 2).

**Figure 2.**
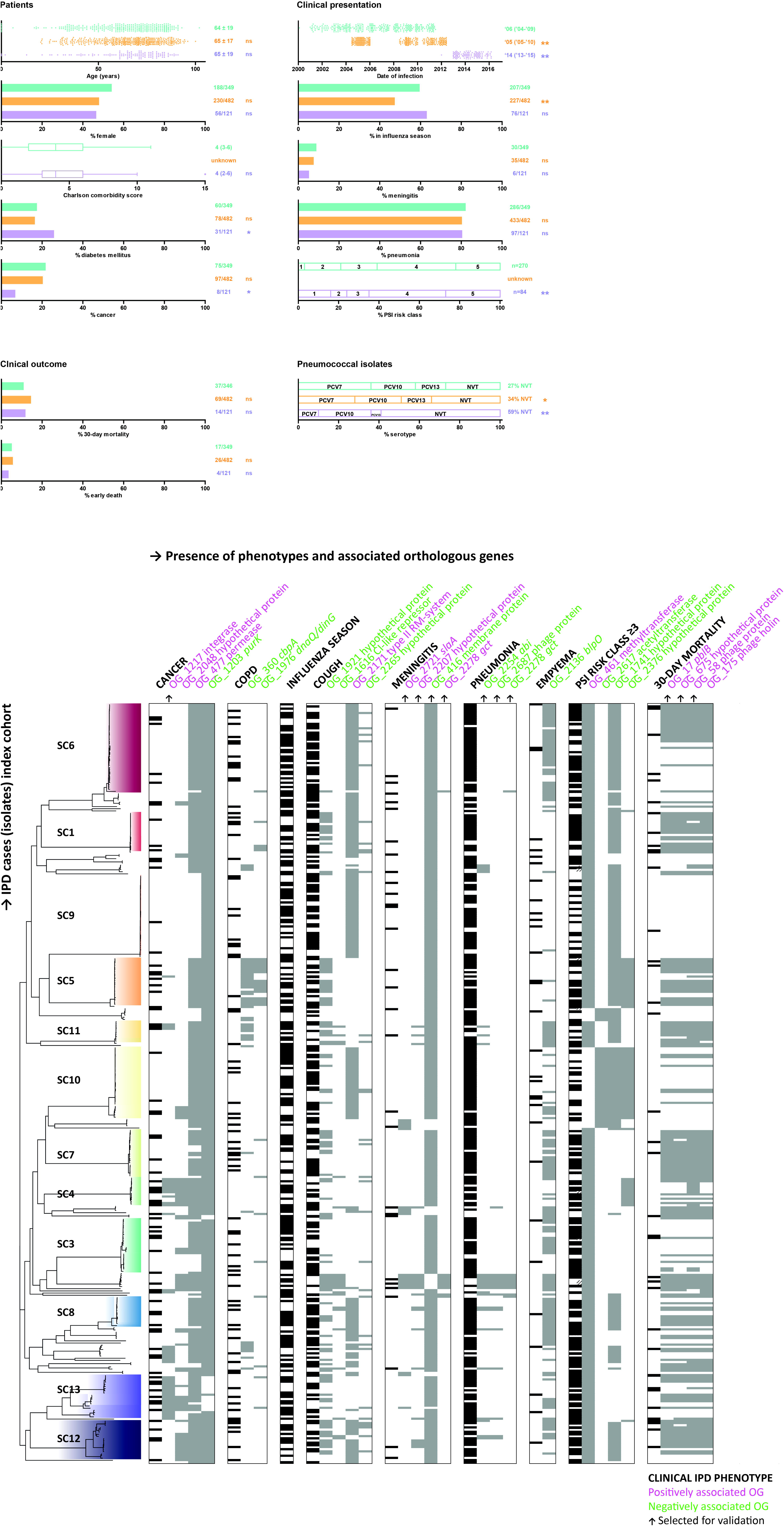
Clinical IPD phenotypes with associated orthologous pneumococcal genes in index cohort. Rows represent 349 IPD cases and corresponding pneumococcal blood isolates. The tree on the left represents their relative phylogenetic position based on SNPs in the core genome, in which sequence clusters are highlighted. The columns represent the presence (filled) or absence (empty) of clinical IPD phenotypes and their associated pneumococcal orthologous genes (OGs) with annotation at the top. Maximally 4 associated OGs that passed Fisher’s exact with p < 0.01 and were independent of population structure are displayed. The OGs selected for validation are indicated by an arrow. *Abbreviations:* IPD: invasive pneumococcal disease; OG: orthologous gene; SC: sequence cluster.

Another method was used to identify genome-wide associations with any allelic variant, or k-mer, present anywhere on the genome and the clinical IPD phenotype. The identified k-mers had nucleotide sequences that aligned with members of up to 6 different OGs. This number of origins was inversely related to k-mer size as well as to the proportion of core (versus accessory) OG origins. None of the 15,249,832 identified unique k-mers met the genome-wide significance threshold in their SC-independent association with clinical IPD phenotypes (Supplementary table 1). Despite this, certain OGs were overrepresented as they contained multiple variable regions related to a particular phenotype (Supplementary table 2).

### Validation of associations

Only associations with OGs were taken into validation, because validation of associations with SCs and k-mers would have required fully sequenced clinical validation cohorts. OG_17 was considered to be a proxy for OG_761 because of their consistency in the index cohort. PCR assays were successful for 8 out of 10 OGs selected for validation (Supplementary methods 3). The size of the temporal validation cohort only allowed for validation of the OGs associated with 30-day mortality. All diluted templates from isolates in both validation cohorts (n=603) were positive for the *gyrA* pneumococcal housekeeping gene with a Ct-value of 24 ± 2. Out of the nine OG-phenotype combinations from the index cohort tested, four were confirmed in the geographical validation cohort (Figure 3) and further characterised as described below.

**Figure 3.**
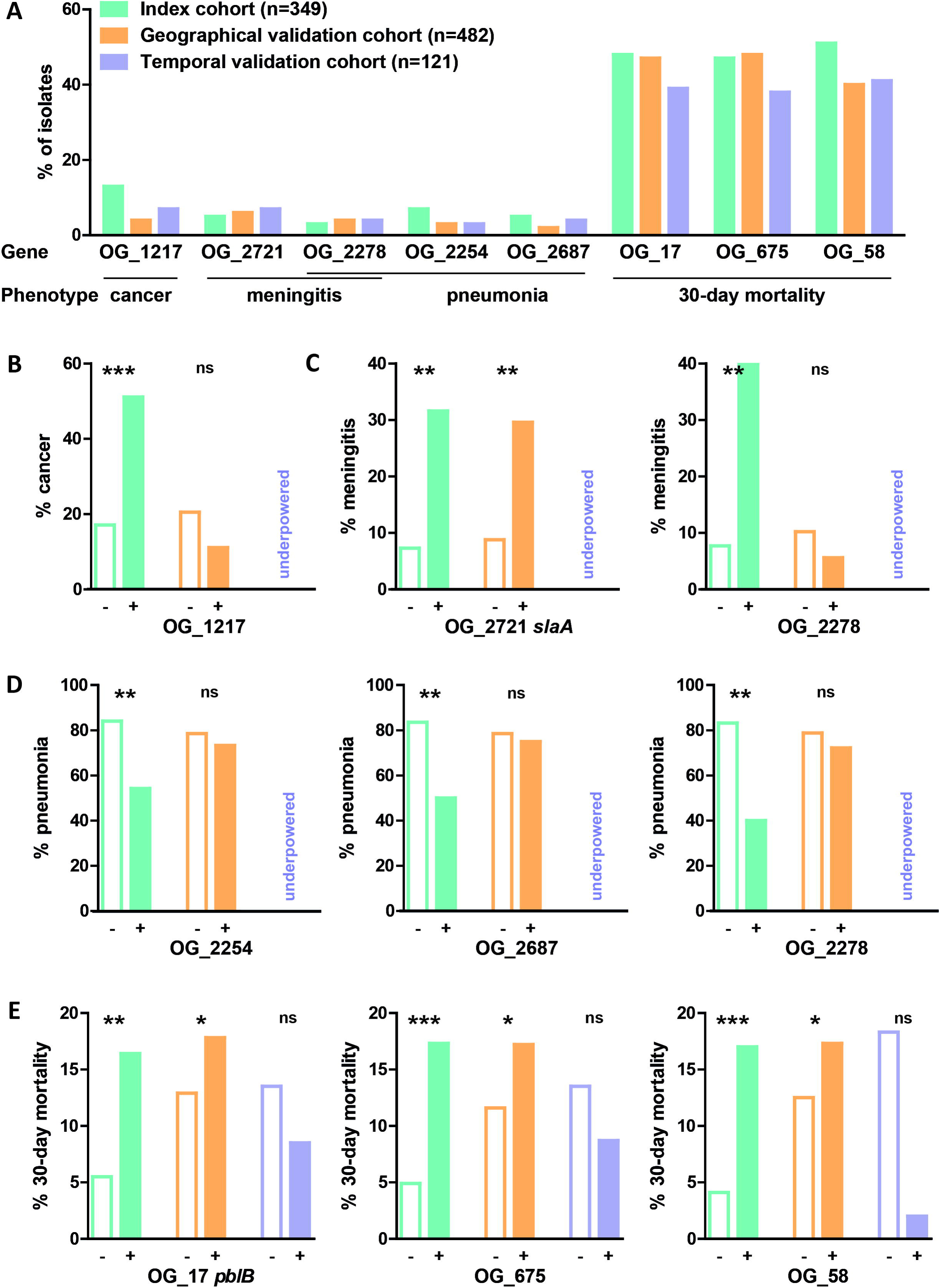
Geographical and temporal validation of orthologous gene associations. The prevalence of pneumococcal OGs in the 3 cohorts (panel A). The association between absence (-, empty bar) or presence (+, filled bar) of and OG and the proportion of patients affected by a particular IPD phenotype in the index and 2 validation cohorts (cancer: panel B; meningitis: panel C; pneumonia: panel D; 30-day mortality: panel E). ^∗^:p<0.1; ^∗∗^:p<0.01; ^∗∗∗^:p<0.001. *Abbreviations:* OG: orthologous gene; IPD: invasive pneumococcal disease. P-values indicate differences in the proportion of patients affected by a particular IPD phenotype.

### Confirmed pneumococcal genotypes

The confirmed OG_2721 related to meningitis was functionally annotated as *slaA* coding for phospholipase A2, and showed 100% anti-occurrence with OG_416 (a predicted membrane protein) and OG_679 (ABC transporter) in the index cohort. The three confirmed OGs related to 30-day mortality were annotated as phage proteins (OG_17 specified as *pblB* encoding a prophage tail fiber protein), and showed high co-occurrence with each other in all three cohorts (Figure 4). In the index cohort, all *pblB* homologues were located either within borders of predicted prophage elements, or located near contig breaks or on short contigs, thus representing circumstances under which prophage elements cannot be identified from draft genomes. While all sequences of OG_2721 were identical, other OGs showed large variation (Supplementary Figure 1). Within OG_17 and OG_58 the number of pairwise amino acid positions exceeded the number expected from their largest sequence variant. This suggests genetic mosaicism which is typical for bacteriophage genes. In the distribution of the confirmed OGs in the pneumococcal populations, OG_17 was taken as a proxy for its joint prophage vector shared with OG_675 and OG_58 (Supplementary Figure 2). In addition to presence, also the number of open reading frames per OG present in an isolate strongly correlated between these three OGs. While all confirmed OGs were present in both vaccine and non-vaccine serotypes, the relative occurrence of OG_2721 in IPD cases remained stable over time, yet OG_17 waned.

**Figure 4.**
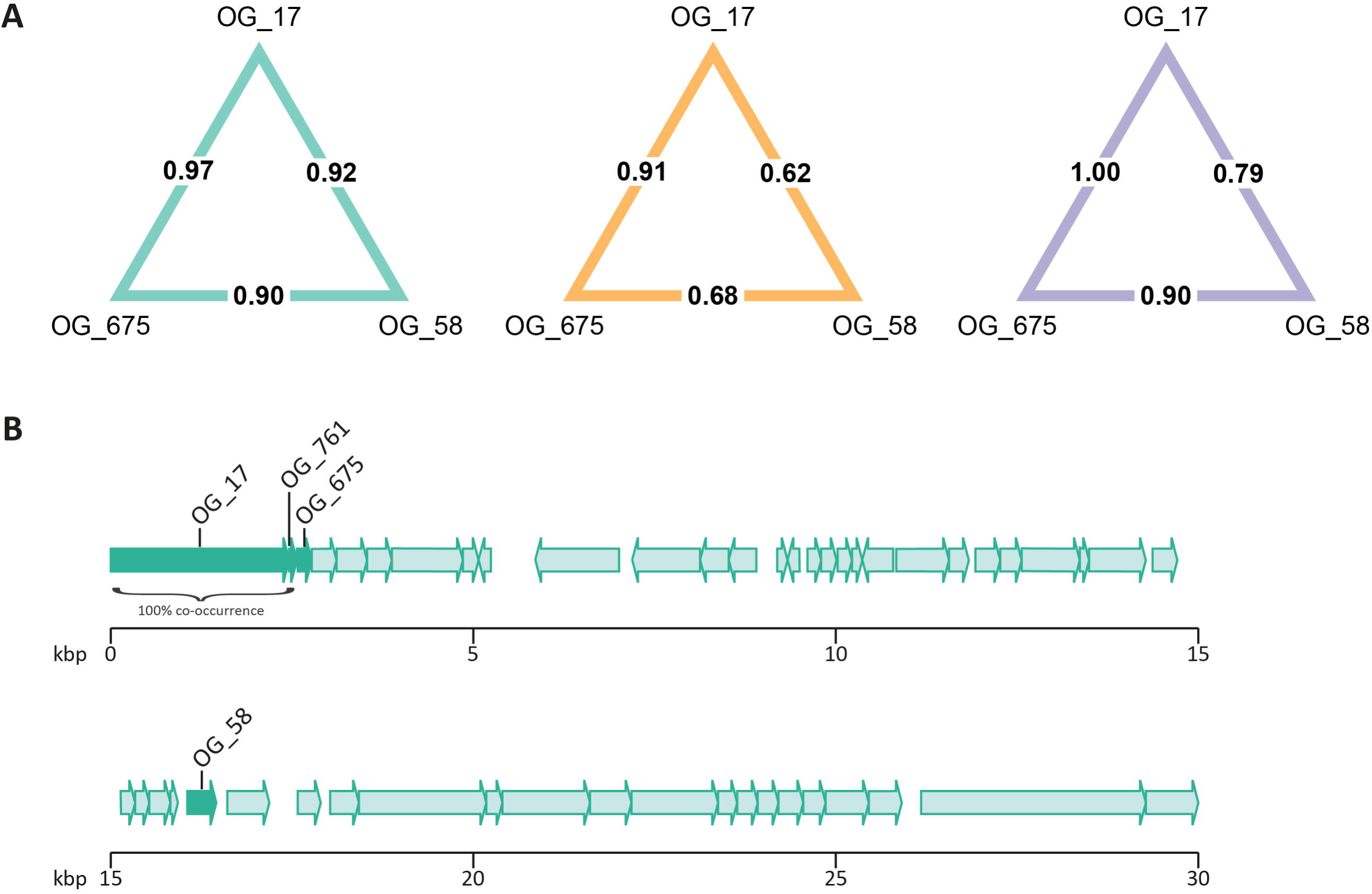
Co-occurrence of mortality-related orthologous genes on single prophage. A. The level of co-occurrence of 3 pneumococcal OGs is expressed in phi coefficient (in bold) for the index, geographical, and temporal cohort (from left to right). B. Gene card depicting an example of the co-localisation of 4 OGs associated with 30-day mortality on a single prophage in pneumococcal isolate PBCN0420. *Abbreviations:* OG: orthologous gene; PBCN: pneumococcal bacteraemia collection Nijmegen; kbp: kilobase pair. P-values support differences from the index cohort.

### Clinical relevance

Relative to clinical predictors of meningitis and 30-day mortality, pneumococcal sequence clusters and orthologous genes were still major independent determinants of these phenotypes (Table 2).

**Table 2.**
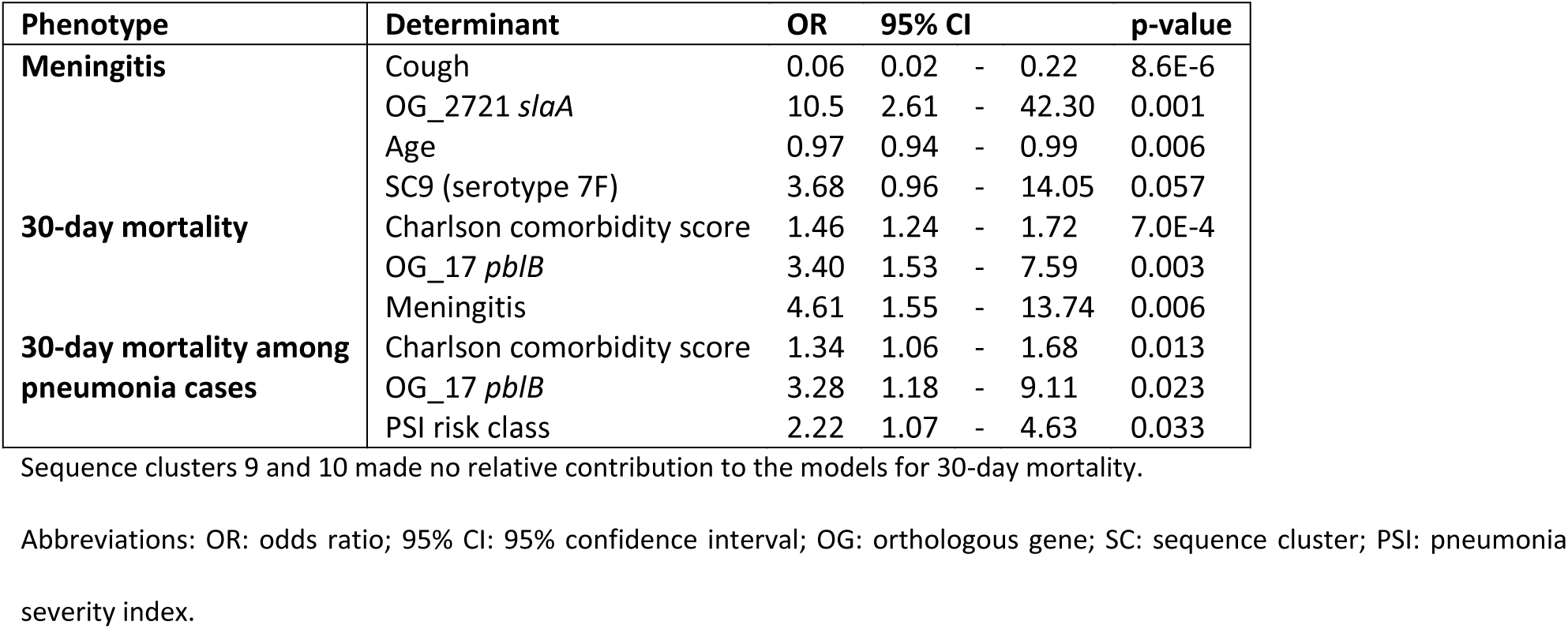
Optimized prediction models for meningitis and 30-day mortality

OG_2721 associated with meningitis was correctly only detected by PCR in serum from patient PBCN0382 (Table 3). For 30-day mortality, OG_675 was most accurately and consistently identified in serum from patients with low pneumococcal DNA loads.

**Table 3.** Detection of orthologous gene sequences in serum from IPD patients

| Study ID |  | PBCN0382 | PBCN0389 | PBCN0420 | PBCN0480 | PBCN0442 |
| --- | --- | --- | --- | --- | --- | --- |
| Meningitis |  | yes | no | no | yes | no |
| 30-day mortality |  | no | yes | yes | yes | no |
| Serum pneumococcal DNA load (copies/ml) |  | 8E+02 | 7E+03 | 3E+03 | 3E+03 | 1E+04 |
| OG_2721<br>*slaA* | Isolate whole genome sequencing <sup>a</sup> | 1 | 0 | 0 | 0 | 0 |
|  | Serum OG-PCR <sup>b</sup> | 2 | 0 | 0 | 0 | 0 |
| OG_17<br>*pblB* | Isolate whole genome sequencing | 2 | 1 | 1 | 1 | 0 |
|  | Serum OG-PCR | 2 | 2 | 1 | 1 | 0 |
| OG_675 | Isolate whole genome sequencing | 2 | 1 | 1 | 1 | 0 |
|  | Serum OG-PCR | 2 | 1 | 2 | 2 | 0 |
| OG_58 | Isolate whole genome sequencing | 2 | 1 | 1 | 1 | 0 |
|  | Serum OG-PCR | 2 | 2 | 2 | 2 | 2 |
| Prophage sequence | Isolate whole genome sequencing | partial | partial | complete | partial | absent |

The results in the matrix represent: <sup>a</sup> number of open reading frames assigned; <sup>b</sup> times target detected in duplicate OG-PCRs.
Abbreviations: IPD: invasive pneumococcal disease; ID: identifier; PBCN: pneumococcal bacteraemia collection Nijmegen; OG: orthologous gene; OG-PCR: orthologous gene polymerase chain reaction.

## Discussion

Through comparative genomics we identified pneumococcal genetic variants (sequence clusters and orthologous genes) to be independent determinants of clinical manifestations of IPD, supported by validation in a separate cohort. These pneumococcal sequence variants could be detected in serum samples from IPD patients by PCR.

Prediction of clinical phenotypes as performed in this study comes with two particular challenges. First, for the identification of certain clinical syndromes one relies on the assessment and examinations performed by the attending physician. Missed diagnosis of for example meningitis cannot be ruled out, given that the absence of cough was one of its main predictors. While uncertainty in sensitivity is inherent to studying clinical phenotypes, the specificity of affected cases is robust as only laboratory confirmed cases of meningitis were classified as such. In fact, instant knowledge of pneumococcal genotype could be used to improve future recognition of particular disease manifestations. Second, although mortality from IPD is more easy to establish, its determinants can vary widely across different clinical settings.^25^ We have observed in our temporal post-vaccination validation cohort, that the relative contribution of pneumococcal variants to mortality may also be influenced by an altered composition of the pneumococcal population itself. Therefore, validity of our findings in other settings should be tested. At the same time, it is difficult to estimate a sample size threshold at which to reject validity because other settings commonly differ in standards of care, population at risk,^26^ antibiotic resistance level, and serotype distribution.^27^ Therefore, although targeted validation as performed in our relatively similar clinical cohorts seems appropriate, in very dissimilar populations *de novo* identification of relevant pneumococcal genotypes may be a more efficient approach.

Our non-selective method including genome-wide pneumococcal variants in naturally occurring IPD populations ensured the likelihood that an association being identified directly correlated with its clinical relevance. In a previous Malawian GWAS where no pneumococcal meningitis-related OGs were identified, not only the human and pneumococcal population differed from ours,^28^ also the heavy selection for meningitis cases could have altered the relative contribution of certain pneumococcal variants.^29^ Vice versa, a determinant identified from an artificial distribution of cases (with unnatural pre-odds), may no longer be valid among patient populations presenting to the hospital.

In general, pneumococcal population structures are characterized by linkage disequilibrium, which means that particular groups of sequences (including the capsular sequence variant) co-occur together on a pneumococcal genome. In our GWAS analyses, to prevent identification of sequences that actually represent a magnitude of co-occurring genes, we corrected for this population structure. Also, we assessed whether these so-called sequence clusters as a whole were related to clinical IPD phenotypes, and we found a remarkable concordance to previous serotype-based studies.^30-32^

Our exclusion of strain- and lineage-specific effects may explain why we have not identified variants of single genes that have previously been described to enhance transition from blood to CSF in laboratory models such as *nanA*, *cbpA*, *pCho*, *lytA*, *ply*, and *glpO*.^33^ At the same time, this aspect of our approach may have favoured the detection of bacterial genotypes located on prophage vectors to be associated with clinical phenotypes as found by us and by others previously.^34^ Unlike many other genes in clonal populations, the distribution of bacteriophages is not as strictly determined by lineage. On the other hand, an important example of a prophage sequence that was associated with the severity of invasive meningococcal disease was discovered by gene-array without bias from correction for population structure.^35^ In any case, if indeed multiple proteins encoded by prophage elements have meaningful interactions with human cells during bloodstream infections, it is more likely that this prophage trait was fixated because of some fitness advantage the bacteriophage or the lysogenic pneumococcus experiences during colonization at the respiratory mucosal surface from where it can actually acquire a viable host.

What we have learned from the k-mer-based GWAS is that the number of lineage-independent allelic variants present in a pneumococcal IPD population is too high for identification of robust associations with particular phenotypes at the current sample size. Although one may have expected OG_17 *pblB*-fragments to be identified in relation to 30-day mortality, the sequences in this orthologous gene were too dispersed to meet the k-mer selection and association thresholds. On the other hand, despite all sequences of OG_2721 being identical, k-mers originating from OG_2721 were still included in the SEER analysis (and positively associated with meningitis), because these k-mers were also represented by a second OG that was more dissimilar and as such made the k-mer meet the selection criteria. These examples demonstrate the complementarity of the two different GWAS methods employed.

While we identified clinical IPD phenotypes to be associated with pneumococcal genes that were independent of pneumococcal lineage (serotype) and clinical predictors, this does not prove causality. We have not included potential confounders like host genotype, a host factor that mediates susceptibility to meningitis^36^ and may simultaneously induce a mucosal environment that welcomes specific pneumococcal variants. On the other hand, evidence for a direct effect of pneumococcal variants in the human bloodstream was demonstrated by measurement of increased activation of human platelets upon interaction with a *pblB*-positive *S. pneumoniae* compared to its knock out variant.^22^ The predicted function of the protein encoded by the *pblB* gene (e.i. functional annotation based on homology) is a phage tail fiber, and its *S. mitis* orthologue was shown to bind to human platelets as well.^37^ Although we have studied pneumococcal blood isolates, it has been shown based on genomic data no adaptation is needed to cross the blood-brain barrier, so DNA sequences from blood isolates seem representative for pneumococci that reach the cerebrospinal fluid and cause meningitis.^38^ Human phospholipase A2, encoded by *slaA*, has been shown to reduce the integrity of the blood-brain barrier *in vitro*, thereby mediating penetration of endothelial cells by group B *Streptococcus*.^39^ In group A *Streptococcus* the presence of *slaA* enhanced the bacterium’s potential for epithelial adherence, colonization and invasive disease.^40^ Also for *slaA* further studies would be required to elucidate its effects *in vivo* and possibilities to avert these during disease. Furthermore, recent advancements in methods to study functional interactions on pneumococcal genomes ^41,42^ may help to improve our understanding of why particular sequence clusters are overrepresented in certain phenotypes.

This study provides evidence that it does matter which pneumococcal variant proliferates in the bloodstream, as it improves our risk assessment in patients affected by IPD. This suggests that the established value of microbial genomics in public health,^43^ outbreak management and combating antimicrobial resistance,^44^ may now be extended to individual patient care. Increased appreciation of eliciting microbial variants, could push the tailored adjunctive measures that are heavily searched for in clinical sepsis care.^45^

Because population dynamics are likely to affect their relative importance, the mapping of microbial variants of concern needs to be supported by strong interdisciplinary surveillance networks. While a systems biology approach may unravel the exact pathophysiology, prompt molecular diagnostics at the emergency department could readily improve risk stratification and alertness for complicated infection in individual patient care.^46^

## Acknowledgements

We thank Dr. A. van der Ende at the Reference Laboratory for Bacterial Meningitis and all hospitals involved in the Dutch national surveillance programme for their concerted efforts which made it possible to validate our findings.

## Supplementary methods

### Supplementary methods 1

Details in handling of clinical variables

#### Data collection

For all 3 cohorts, clinical data were collected from medical charts and registered with patient identifiers in a secured source file. In a separate working file, labeled clinical data were stored together with the non-identifying study code assigned to each IPD case included. The following clinical data were only collected for patients admitted to the index hospitals: Charlson comorbidity score, COPD, cardiovascular disease, antibiotics prior to admission, SIRS, blood CRP and leukocyte count, PSI risk class, and time to death. Otherwise, clinical data were handled in an equal manner across the 3 cohorts.

#### Variables

The following clinical variables were defined, processed or classified in a particular way. Included invasive pneumococcal disease cases: *S. pneumoniae* isolated from culture of cerebrospinal fluid or blood. Charlson comorbidity score: calculated for cases ≥18 years old. Immunocompromising therapy: actual use of systemic corticosteroids or chemotherapy. Cardiovascular disease: history of either hypertension, myocardial infarction, myocardial insufficiency, claudicatio intermittens, vasculitis, vascular stents, heart catheterization, atrial fibrillation or hypercholesterolemia. Antibiotics prior to admission: within preceding week either in context of separate medical issue or for current infection. Date of infection: date of blood culture collection. Influenza season: defined annually from the first to the last week with >5 reported influenza cases in the Netherlands as reported by WHO FluNet (http://apps.who.int/flumart/Default?ReportNo=12). Systemic inflammatory response syndrome: percentage of immature neutrophils not accounted for. Cough: as reported or observed at admission. Pneumonia, empyema, and meningitis were not mutually exclusive. Pneumonia, meningitis, and unknown focus of infection: as reported by the attending physician. Pleural effusion: as reported by the attending radiologist. Empyema: *S. pneumoniae* isolated from pleural fluid culture. CRP, leukocytes, and other chemistry results included in clinical algorithms: at day of admission, except for bilirubin and albumin during hospital stay. Pneumonia severity index score: calculated for cases ≥18 years old who suffered from pneumonia, formatted into PSI risk class and stratified to PSI risk class 1-2 versus 3-5. 30-day mortality: in hospital death within 30 days from admission. Early death: in hospital death within 48 hours from admission.

#### Missing data

Missing data were not replaced. Data were considered to be missing if the corresponding section was not present in the medical chart. PSI risk class was only considered valid and reported if ≥16 included variables were known. All other clinical algorithms were reported only if missing variables did not influence case classification.

### Supplementary methods 2 Interpretation of real time PCR amplification and melting curves

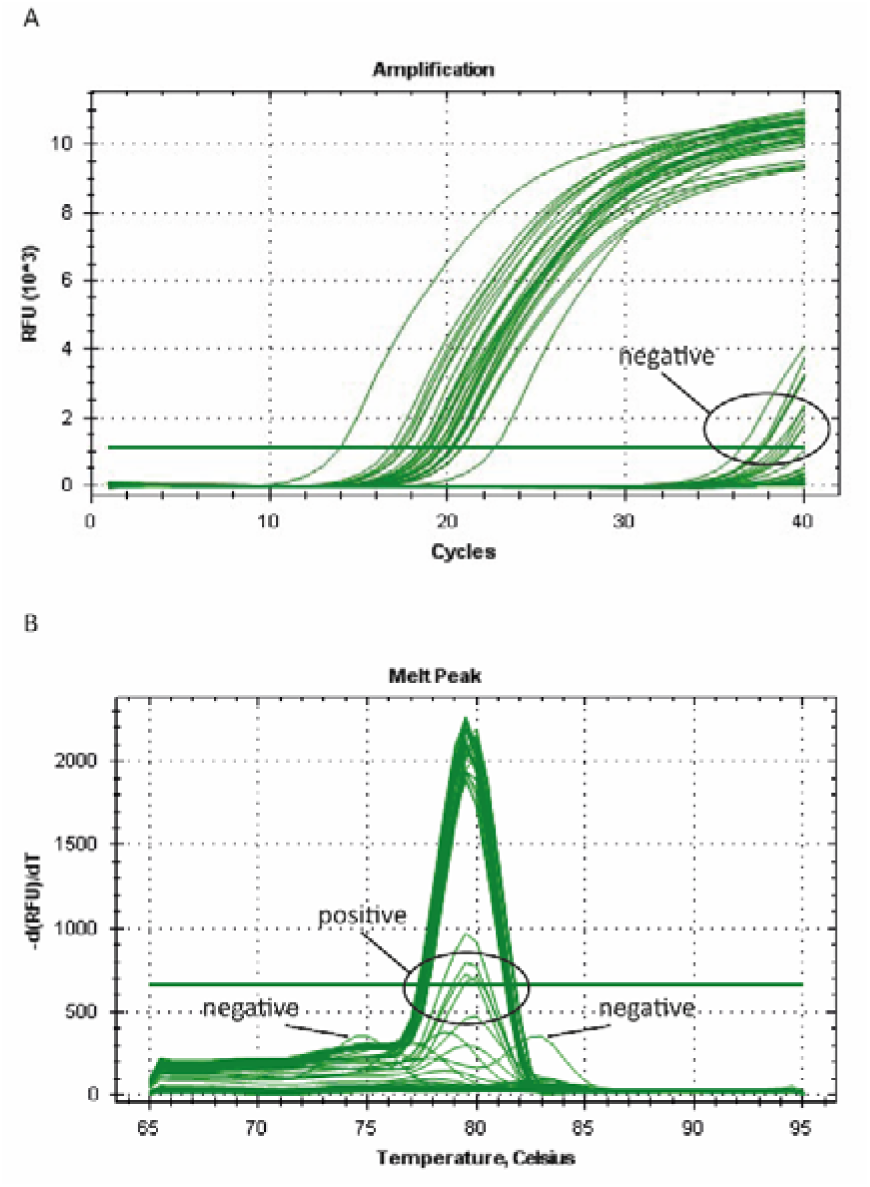

### Supplementary methods 3 Primer characteristics

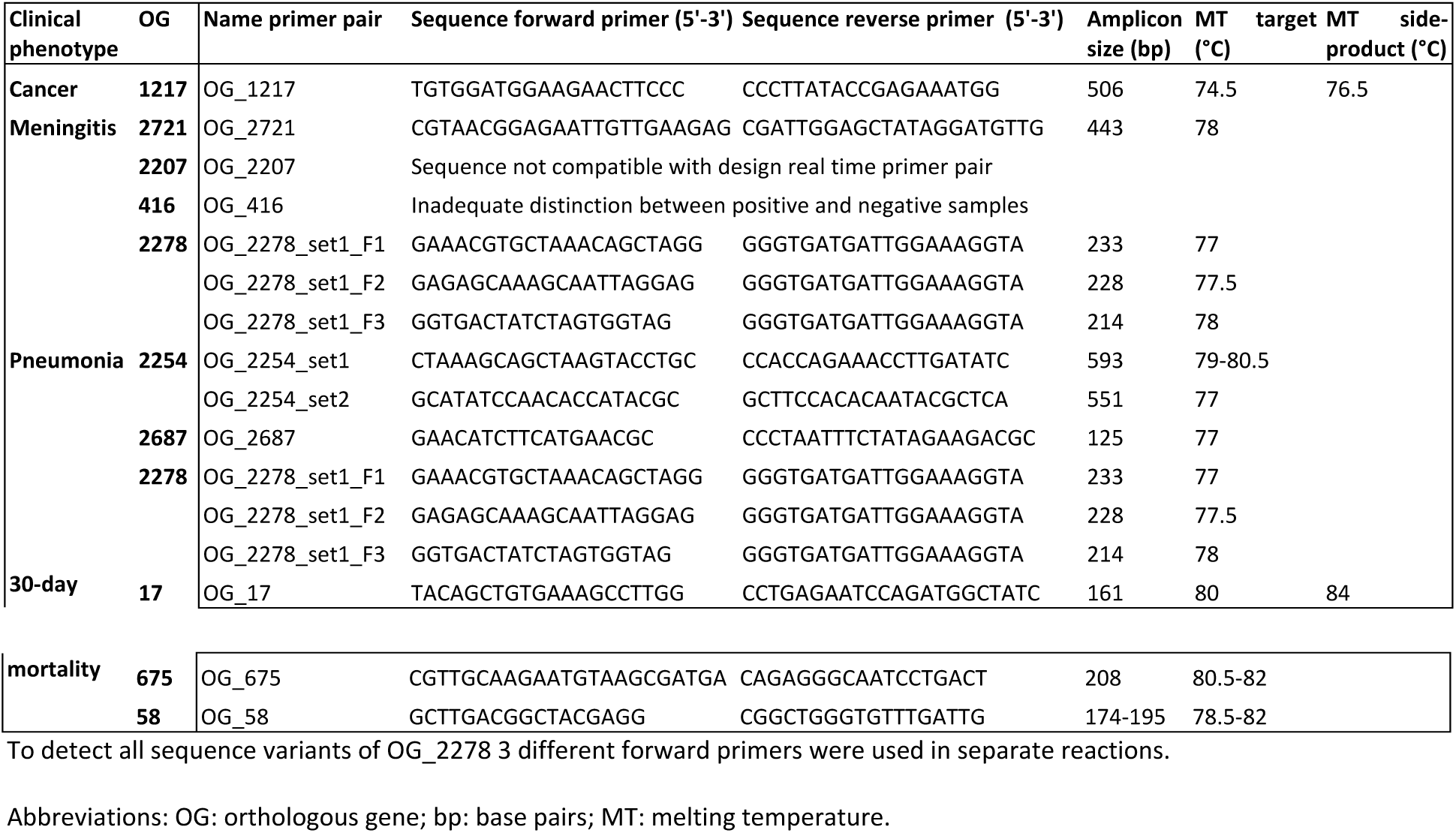

**Supplementary figure 1.**
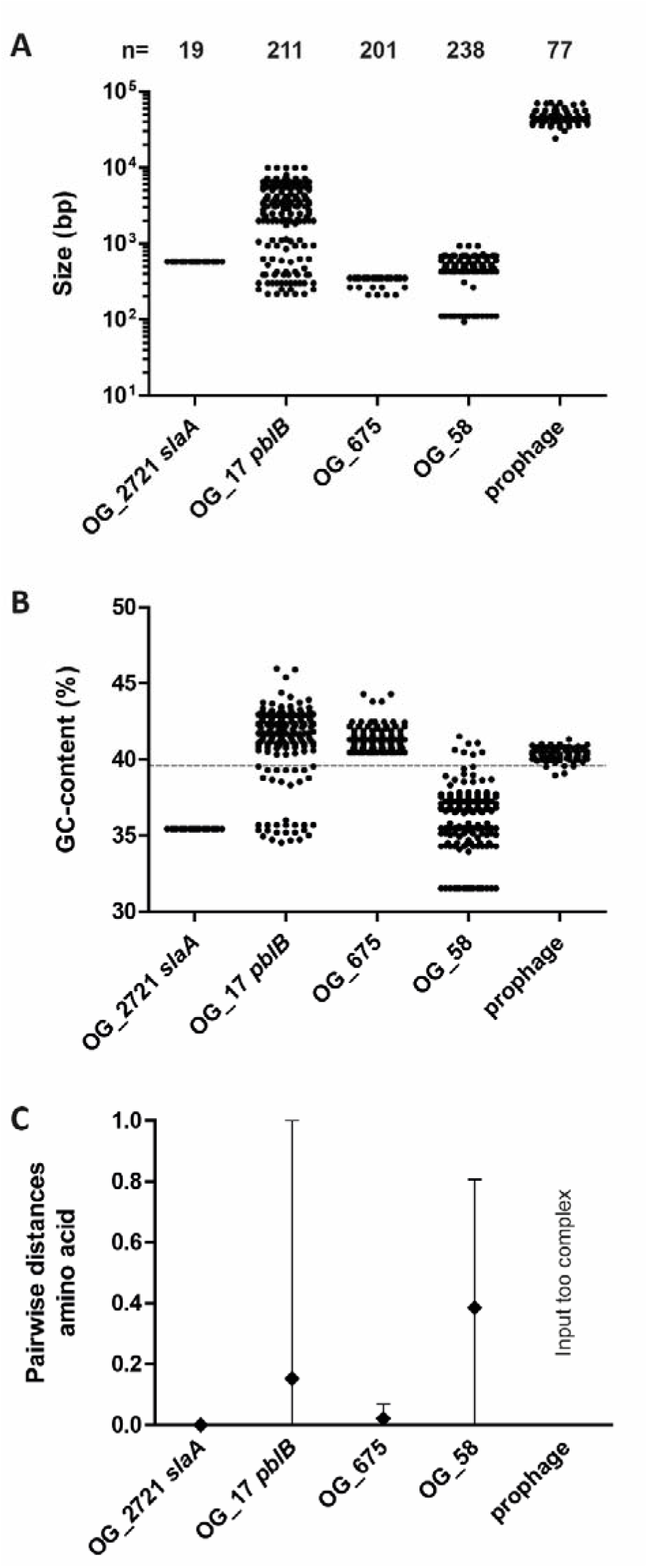
Sequence variation within confirmed orthologous genes

**Supplementary figure 2.**
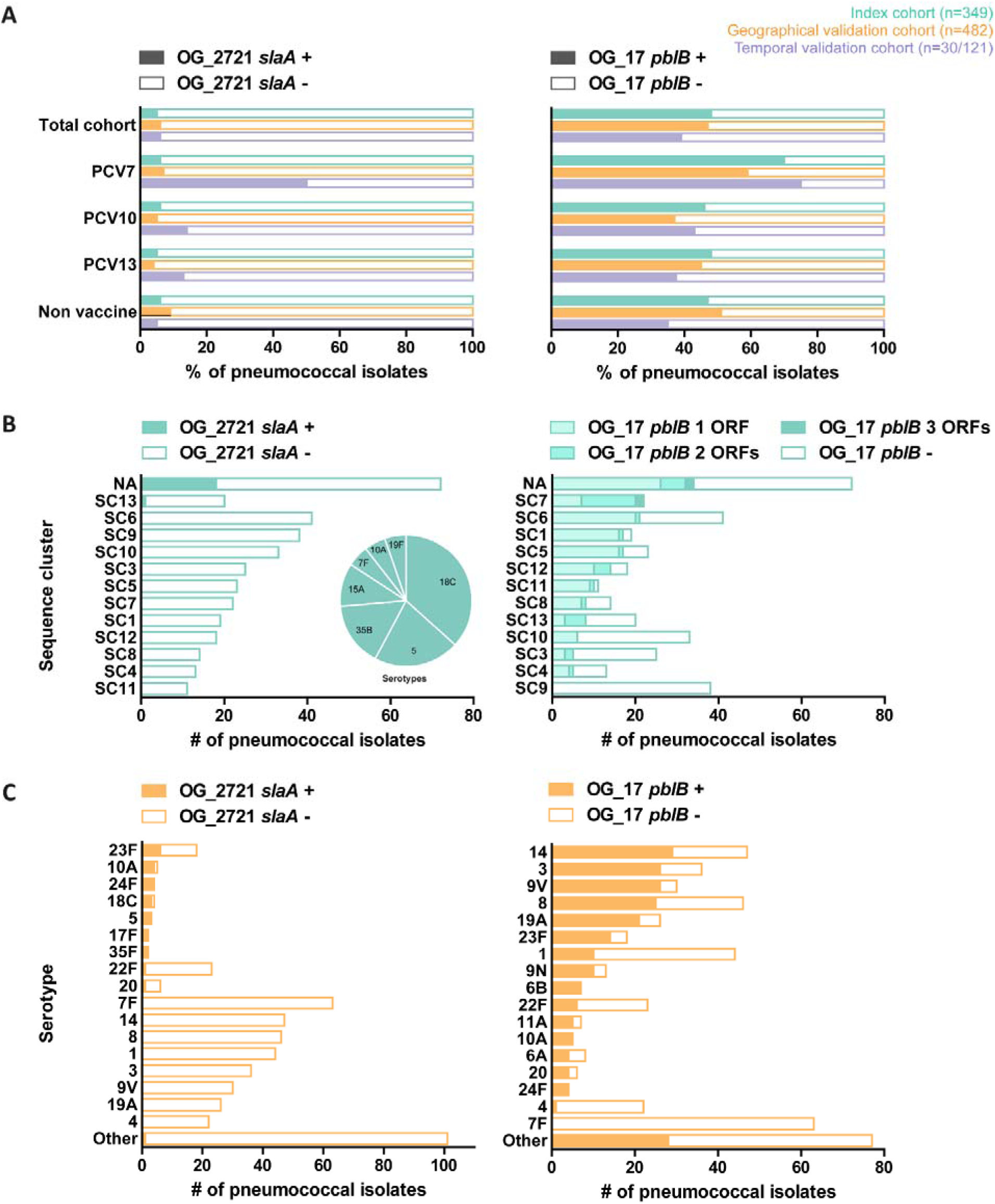
Distribution of 2 confirmed orthologous genes in the pneumococcal population

**Supplementary table 1.** Individual k-mers associated with clinical IPD phenotype in index cohort p<e-6

| Phenotype | K-mer | OG | Core OG | Annotation | Direction | p-value |
| --- | --- | --- | --- | --- | --- | --- |
| **Pneumonia** | X10_9 | 856 | no | gp19 | neg | 7E-07 |
|  | X10_9 | 163 | no | zinc metalloproteinase | neg | 7E-07 |
| **Meningitis** | X11_41 | 2093 | no | transmembrane protein | pos | 6E-07 |
|  | X11_32 | 340 | yes | DNA polymerase III subunits gamma and tau | neg | 7E-07 |
|  | X11_86 | 10 | no | ABC transporter ATP-binding protein/permease | pos | 9E-07 |
| **Immunocompromising therapy** | X3_1 | 35 | no | endo-beta-N-acetylglucosaminidase | neg | 3E-07 |
|  | X3_1 | 212 | no | ABC transporter ATP-binding protein/permease | neg | 3E-07 |
|  | X3_1 | 175 | no | phage holin | neg | 3E-07 |
|  | X3_1 | 1078 | yes | phosphatase | neg | 3E-07 |
| **Diabetes mellitus** | X5_1 | 1415 | yes | 6-phosphogluconolactonase | neg | 8E-08 |
|  | X5_5 | 521 | yes | ABC transporter permease | neg | 8E-08 |
| **Antibiotics prior to admission** | X8_1 | 870 | no | UDP-glucuronate 5'-epimerase | neg | 5E-09 |
|  | X8_1 | 255 | yes | excinuclease ABC subunit A | neg | 5E-09 |
|  | X8_1 | 1277 | yes | flavoprotein | neg | 5E-09 |
|  | X8_10 | 323 | yes | alpha-amylase | neg | 5E-09 |
|  | X8_2 | 335 | no | metallo-beta-lactamase superfamily protein | neg | 5E-09 |
|  | X8_2 | 169 | no | hypothetical protein | neg | 5E-09 |
|  | X8_3 | 1701 | no | BlpM | neg | 5E-09 |
|  | X8_4 | 216 | yes | DNA polymerase I - 3'-5' exonuclease and polymerase | neg | 5E-09 |
|  | X8_5 | 967 | yes | hypothetical protein | neg | 5E-09 |
|  | X8_5 | 610 | no | serine/threonine protein kinase | neg | 5E-09 |
|  | X8_5 | 243 | no | HAD superfamily hydrolase | neg | 5E-09 |
|  | X8_7 | 546 | no | single-stranded DNA-binding protein | neg | 5E-09 |
|  | X8_7 | 3012 | no | hypothetical protein | neg | 5E-09 |
|  | X8_14 | 192 | yes | single-stranded DNA-specific exonuclease | pos | 5E-09 |
|  | X8_14 | 1533 | no | UDP-N-acetylglucosamine-2-epimerase | pos | 5E-09 |
|  | X8_15 | 6 | no | cell-division ATP-binding protein FtsE | pos | 5E-09 |
|  | X8_16 | 1266 | yes | GIY-YIG domain-containing protein | pos | 5E-09 |
|  | X8_17 | 321 | yes | cation transporter E1-E2 family ATPase | pos | 5E-09 |
|  | X8_18 | 2968 | no | GNAT family acetyltransferase | pos | 5E-09 |
|  | X8_18 | 221 | no | tyrosine-protein phosphatase CpsB | pos | 5E-09 |
|  | X8_19 | 932 | yes | xanthine phosphoribosyltransferase | pos | 5E-09 |
|  | X8_19 | 670 | no | lactoylglutathione lyase | pos | 5E-09 |
|  | X8_20 | 1359 | no | dTDP-4-dehydrorhamnose 3 | pos | 5E-09 |
|  | X8_21 | 26 | no | oligopeptide binding lipoprotein | pos | 5E-09 |
|  | X8_22 | 12 | no | IS1239 transposase | pos | 5E-09 |
|  | X8_22 | 1232 | no | branched-chain aa ABC transporter permease | pos | 5E-09 |
|  | X8_23 | 347 | yes | Gfo/Idh/MocA family oxidoreductase | pos | 5E-09 |
|  | X8_23 | 10 | no | ABC transporter ATP-binding protein/permease | pos | 5E-09 |
|  | X8_26 | 21 | no | cell envelope integrity inner membrane protein Tol | pos | 5E-09 |
|  | X8_29 | 669 | yes | dihydrofolate reductase | pos | 5E-09 |

**Supplementary table 2.**
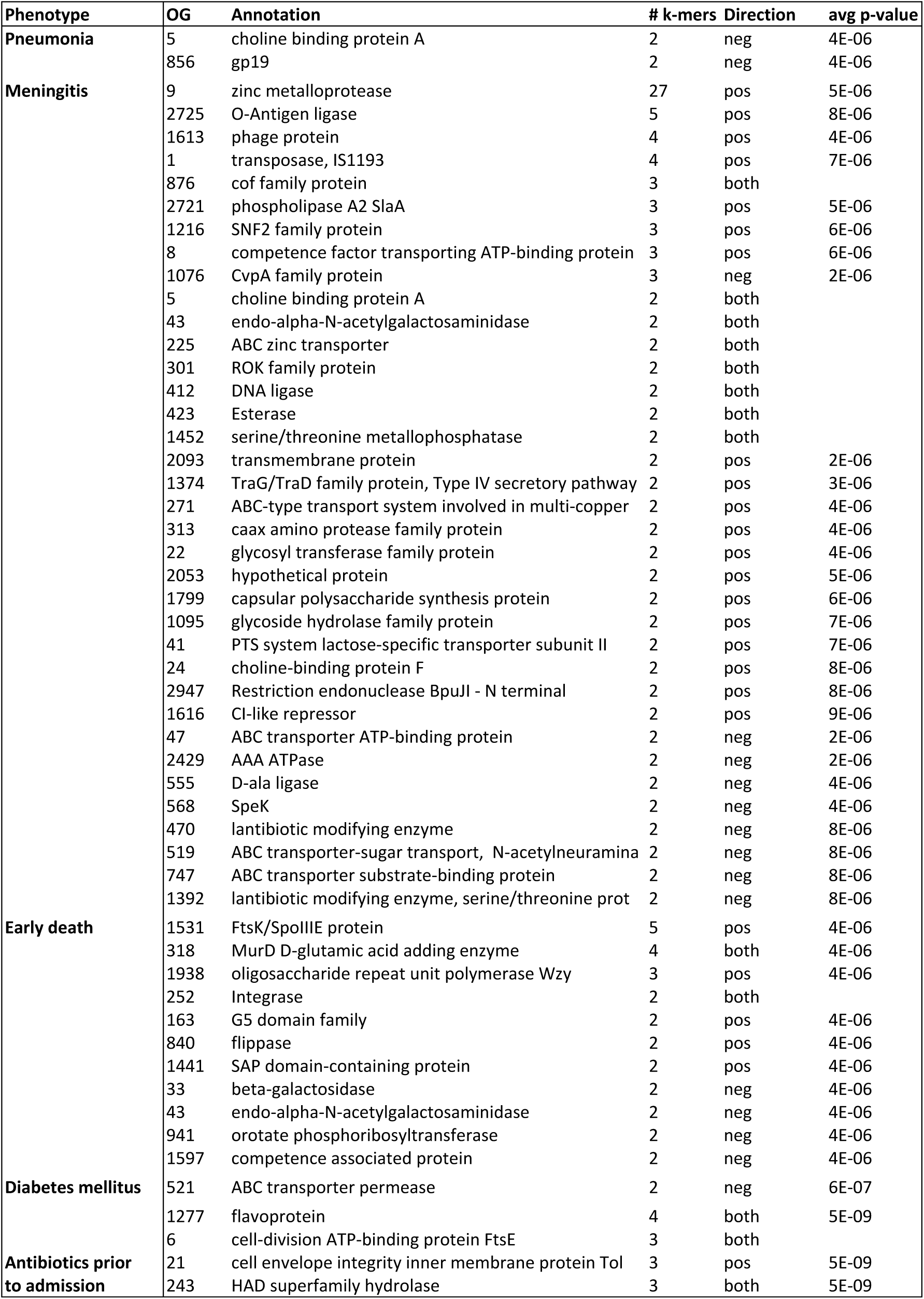
OG‐clustered k‐mers associated with clinical IPD phenotype in index cohort

